# Establishment of epithelial and fibroblast cell lines from primary renal cancer nephrectomies

**DOI:** 10.1101/337055

**Authors:** Ning Yi Yap, Teng Aik Ong, Christudas Morais, Jayalakshmi Pailoor, Glenda C. Gobe, Retnagowri Rajandram

## Abstract

Renal cell carcinoma (RCC) is one of the most lethal urogenital cancers and effective treatment of metastatic RCC remains an elusive target. Cell lines enable the in-vitro investigation of molecular and genetic changes leading to renal carcinogenesis and are important for evaluating cellular drug response or toxicity. This study details a fast and easy protocol of establishing epithelial and fibroblast cell lines concurrently from renal cancer nephrectomy tissue. The protocol involves mechanical disaggregation, collagenase digestion and cell sieving for establishing epithelial cells while fibroblast cells were grown from explants. This protocol has been modified from previous published reports with additional antibiotics and washing steps added to eliminate microbial contamination from the surgical source. Cell characterization was carried out using immunofluorescence and quantitative PCR. Eleven stable epithelial renal tumour cell lines of various subtypes, including rare subtypes, were established with a spontaneous immortalization rate of 21.6% using this protocol. Eight fibroblast cell cultures grew successfully but did not achieve spontaneous immortalization. Cells of epithelial origin expressed higher expression of epithelial markers such as pan-cytokeratin, CK8 and E-cadherin whereas fibroblast cells expressed high α-SMA. Further mutational analysis is needed to evaluate the genetic or molecular characteristics of the cell lines.

## Introduction

Renal cell carcinoma (RCC) comprises of 2-3% of all human malignancies and the incidence is increasing worldwide [1]. The incidence of RCC is higher in Western countries (North America, Europe, Australia and New Zealand) compared to Asian countries in general [1]. The most common subtypes of RCC are clear cell (70-80%) followed by papillary (10%), chromophobe (5%) and collecting duct RCC (1%) [2]. RCC with sarcomatoid or rhabdoid transformation is not a recognised subtype of RCC as sarcomatoid or rhabodoid features can be found in all histologic subtypes of RCC [3]. Both of these histological transformations in RCC are associated with aggressive tumours and poor prognosis. Radiation and chemotherapy has limited efficacy on RCC. The current approved targeted therapies for metastatic RCC suppress angiogenesis by inhibiting vascular endothelial growth factor (VEGF) or platelet-derived growth factor (PDGF) mediated pathways. However, most patients eventually develop resistance to these drugs and require second or third-line therapies [4]. Immune checkpoint inhibitors are the most recent approved for treatment of metastatic RCC. Nevertheless, since these drugs are relatively new, the long term clinical outcome is unknown at the moment. Hence, the genetic and molecular changes leading to the pathogenesis of RCC and development of therapy resistance are still not well understood.

Cell lines provide an avenue for in-vitro investigation into the molecular and genetic aspects of renal cancer carcinogenesis and pre-clinical studies for evaluating drug response or toxicity at a cellular level. Primary cell cultures offer the advantage of being more biologically similar to the tumour while continuous cell lines are homogenous and could be sub-cultured indefinitely. Both have their advantages and are integral for in-vitro experimentation. At present, most commercially available RCC cell lines are established from Caucasians, such as ACHN, A-498, Caki-1, Caki-2, 769-P and 786-O. Asians and Caucasians differ in the incidence of RCC and their response to targeted treatment and immunotherapy [1,5,6]. Hence, there might be some difference in the underlying molecular mechanisms of RCC cells in Caucasians and Asians. The advantage of establishing cell lines from any research centre’s population is that patient information at diagnosis, tumour aggressiveness or patient clinical outcome of the corresponding cell lines will be known. In addition, the molecular responses of the established cell lines will likely conform to the characteristics of the study population.

In this study, a fast and simple method of establishing RCC epithelial and fibroblast as well as normal kidney epithelial cortex cell lines from primary tumours or nephrectomy specimen collected at surgery is described, with emphasis on RCC epithelial cell lines. The outcome of the cell line establishment and methods of initial cell characterization are also presented here. This protocol has been modified from previous published reports with additional antibiotics (Primocin and Mycozap) and washing steps added to eliminate yeast/mold, bacterial and mycoplasma contamination present from the surgical source [7-13]. In addition, this is the first report to present the details and protocols of simultaneous establishment of RCC, normal kidney and RCC associated fibroblast cell lines from multiple trials/tumours.

## Methods and materials

RCC tissue and normal kidney samples were collected from consented patients who have undergone nephrectomy for the removal of kidney tumour. Ethical approval was obtained from the University of Malaya Medical Centre (UMMC) Ethics Committee (Ref : 848.17) and written informed consent was obtained for each patient.

### Materials for cell line establishment

Table 1 lists out the solutions or media which were required for the establishment of cell lines. Unless stated otherwise, high glucose Dulbecco’s Modified Eagle’s Medium (DMEM) was purchased from Nacalai Tesque (Japan) and cell culture materials or supplements were obtained from Gibco (USA).

**Table 1.**
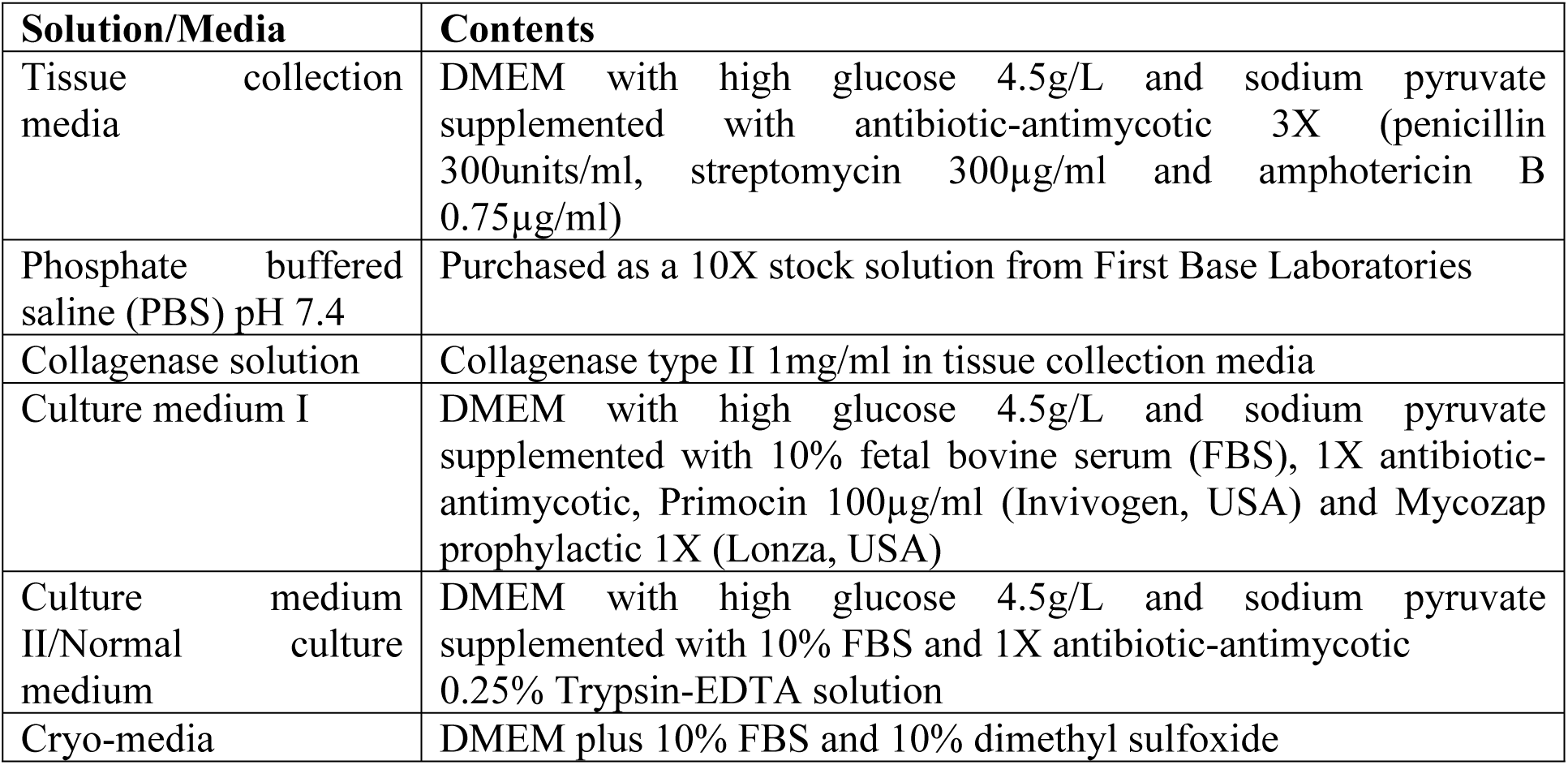
Solutions or media used in the establishment and maintenance of cell lines

| **Solution/Media** | **Contents** |
| --- | --- |
| Tissue collection media | DMEM with high glucose 4.5g/L and sodium pyruvate supplemented with antibiotic-antimycotic 3X (penicillin 300units/ml, streptomycin 300µg/ml and amphotericin B 0.75µg/ml) |
| Phosphate buffered saline (PBS) pH 7.4 | Purchased as a 10X stock solution from First Base Laboratories |
| Collagenase solution | Collagenase type II 1mg/ml in tissue collection media |
| Culture medium I | DMEM with high glucose 4.5g/L and sodium pyruvate supplemented with 10% fetal bovine serum (FBS), 1X antibiotic-antimycotic, Primocin 100µg/ml (Invivogen, USA) and Mycozap prophylactic 1X (Lonza, USA) |
| Culture medium II/Normal culture medium | DMEM with high glucose 4.5g/L and sodium pyruvate supplemented with 10% FBS and 1X antibiotic-antimycotic 0.25% Trypsin-EDTA solution |
| Cryo-media | DMEM plus 10% FBS and 10% dimethyl sulfoxide |

### Tissue collection

Tissue collection was carried out aseptically with the help of the urologist surgeon and pathologist. The urologist surgeon confirmed the location of the RCC lesions during tissue collection and the pathologist confirmed the pathological diagnosis after tissue processing. RCC tissue samples were taken within the tumour region, away from the tumour margin. Tumour samples were collected from the fleshy soft part of the tumour, avoiding necrotic, fibrotic or haemorrhagic areas. Normal kidney samples were collected from the outer kidney cortex with macroscopically normal appearance which was furthest away from the tumour lesion. Tissue collection was carried out within an hour of the tumour or kidney removal from the patient. Fig 1 illustrates an example of a resected kidney with RCC from which tumour tissue was taken for cell line establishment. Each tissue sample was cut in two pieces with one part for formalin fixation (formalin fixed paraffin embedded/FFPE slides) and one part for cell line establishment. The RCC tissues for cell line establishment (sized 0.5-1.5cm^3^) were placed in separate 50ml centrifuge tubes with 5ml of ice-cold tissue collection media.

**Fig 1.**
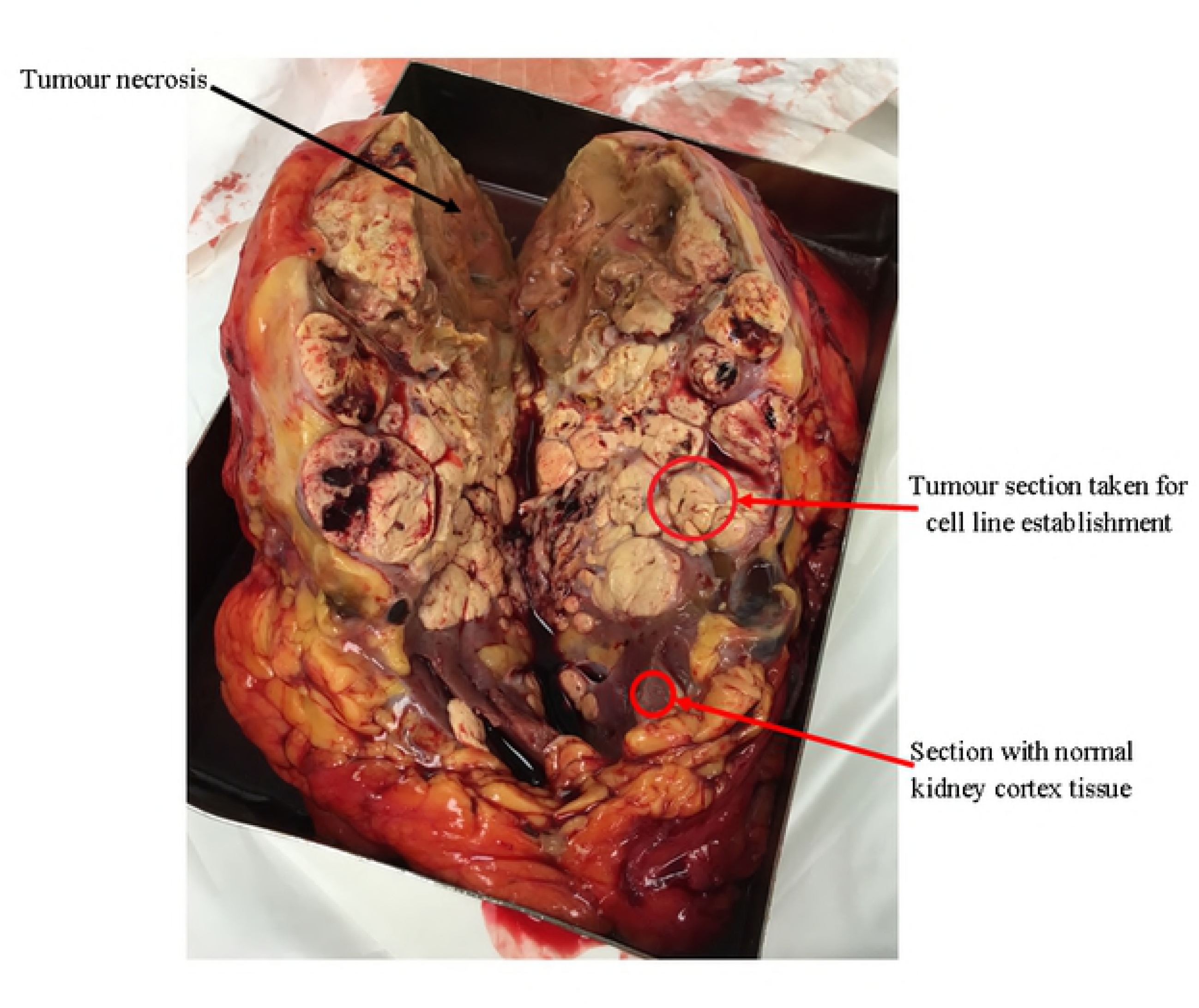
An example of nephrectomy specimen taken before resection of tissues for cell line establishment. The tumour tissue for cell line establishment was taken from a tumour area clear of necrosis or haemorrhage. The section with normal kidney tissue was distinguishable from the tumour lesion.

### Tissue Processing

Tissue samples were transported to the laboratory within an hour after collection and tissue processing was performed aseptically in a Class II biosafety cabinet. Kidney tumour and normal tissue were processed separately using a similar protocol. Tissue samples were first placed on a petri dish where fat tissue and visible blood clots were dissected out. Tissue samples were then washed and agitated in cold PBS pH7.4 in sterile 50ml tubes 4-5 times to remove any remaining blood. At this point, it is possible to store the tissue in culture medium I at 4°C overnight (≈ 8-12 hours) and continue processing the next day. However, it is advisable to proceed to the next steps immediately if time allows.

Tissue samples were placed on 60mm petri dishes and minced into 1mm^3^ pieces with sterile blades. Next, tissue for establishment of epithelial cells was digested with collagenase (Establishment of epithelial cell lines). However, for fibroblast cell establishment, the minced pieces were washed in PBS and placed directly in culture flasks for propagation, without collagenase digestion (Establishment of fibroblast cell lines).

### Establishment of tumour and normal epithelial cell lines

Procedures were performed at room temperature (≈27°C) unless stated otherwise. Tumour and normal epithelial cells were processed using a similar protocol. Tissue fragments were transferred to clean 50ml tubes and washed twice in fresh cold tissue collection media by centrifuging at 300g for 5 minutes. The supernatant was removed each time, and after the second wash, approximately 5ml of collagenase solution was added to each tube. The tubes containing the tissue fragments were agitated using a shaking incubator at 37°C for 45 minutes-1 hour. After enzymatic digestion, the digested tissue from each sample was sieved through a 70μm cell strainer (SPL Life Sciences, South Korea) into a clean 50ml tube to remove undigested tissue and glomeruli. The 50ml tubes containing the cells were centrifuged for 5 minutes at 300g and the supernatant was pipetted off. The cells were re-suspended in pre-warmed culture medium I and transferred to 25cm^2^ culture flasks (SPL Life Sciences, South Korea). Typically, cells from each sample were seeded in two 25cm^2^ flasks. Cell viability was not determined at this stage and seeding density was not tightly controlled. Cells are incubated at 37°C in a humidified atmosphere of 5% CO_2_. After an overnight incubation (12-24 hours), the culture medium in the flasks was pipetted off along with any unattached cells and blood cells. Attached cells were gently washed once with pre-warmed wash media and replaced with fresh culture medium I. This procedure was repeated after 48 and 72 hours until the flasks were free of unattached cells and debris. Subsequently, culture medium was changed every 2-3 days until confluency. On day 7 onwards, culture medium II was used instead of culture medium I.

Yeast and bacterial contamination were tested using the Cell Culture Contamination Detection Kit (Molecular Probes, Thermo Fisher, USA) and presence of mycoplasma was detected using the MycoFluor™ Mycoplasma Detection Kit (Molecular Probes, Thermo Fisher, USA).

### Establishment of cancer associated fibroblast cell lines

Fibroblast cells were grown using the tissue explant technique. Tumour tissue minced into 1mm^3^ pieces was placed in a 25cm^2^ culture flask and pre-warmed culture medium I was added before incubation at 37°C in a humidified atmosphere of 5% CO_2_. After an overnight incubation (12-24 hours), the culture medium in the flasks was pipetted off along with any unattached cells and debris. Attached tissue pieces were gently washed once with pre-warmed wash media and replaced with fresh culture medium I. This procedure was repeated after 48 and 72 hours until the flasks were free of unattached cells and debris. On day 7 onwards, culture medium II was used instead of culture medium I. Fibroblast cells typically migrate out from the explant after 5-10 days. If growth was slow, 5ng/ml of fibroblast growth factor (FGF) (Merck Millipore, USA) was added to the culture medium to encourage fibroblast growth.

### Cell culture maintenance and subculture

Cells were grown to 80-90% confluency before passaging. Culture medium was removed and 1ml of 0.25% trypsin-EDTA was added to each 25cm^2^ flask. After 5 minutes incubation at 37°C, the flasks were gently tapped to detach cells and trypsin reaction was stopped with the addition of 1ml culture medium II. Cells were pelleted by centrifuging at 200g for 5 minutes and re-suspended in culture medium II before seeding into a new 25cm^2^ flask. Passaging was carried out in a 1/2 split ratio. For cryopreservation, cells were re-suspended in 1ml cryomedia, frozen at −80°C overnight and stored at −190°C in a liquid nitrogen tank. Cells were reactivated from cryopreservation by thawing at 37°C in a water bath and centrifuging the cells at 200g for 5 min. The cells were re-suspended in culture medium II and seeded into a 25cm^2^ flask.

### Cell characterization by immunofluorescence staining

General cell morphology was viewed under an inverted microscope. Cells were seeded at 1×10^5^/well in a 24 well plate. One sterile 9mm cell culture coverslip (SPL, South Korea) was placed in each well before seeding. After 24 hours, the growth medium was removed and cover slips were washed 2x with PBS. Cells grown on coverslips were fixed with 4% paraformaldehyde in PBS for 20 minutes at room temperature (RT). The cells were then washed with PBS and permeabilized with 0.1% Triton X-100 in PBS for 10 minutes. Endogenous peroxidase activity was blocked with 3% hydrogen peroxide (H_2_O_2_) in PBS for 5 minutes. After washing with PBS, the cells were incubated with primary antibody diluted to the optimised concentration in PBS overnight at 4°C. The dilution for pan-cytokeratin (PCK-26)(Abcam, USA) was 1:150, alpha smooth muscle actin (α-SMA) (1A4)(Dako, USA) was 1:800, aquaporin-1 (AQP-1) (B-11)(Santa Cruz, USA) was 1:150 and Tamm–Horsfall protein (THP) (B-2) (Santa Cruz, USA) was 1:200. The cells were washed with PBS and incubated with secondary antibody (anti-mouse/rabbit Alexa Fluor 488 from Thermo Fisher) (1:200 dilution) before cover slips were mounted on slides using Vectashield aqueous mounting medium with DAPI (4’,6-diamidino-2-phenylindole).

Immunohistochemistry staining was also carried out using pan-cytokeratin (Abcam, USA)(1:200 dilution), α-SMA (Dako, USA)(1:400 dilution), AQP-1 (Santa Cruz, USA)(1:150) and THP (Santa Cruz, USA)(1:150) in FFPE normal kidney and ccRCC tissue collected from the operating theatre (results not shown). This was done to determine the localisation and expression pattern of both antibodies in intact kidney tissue.

### Quantitative PCR (qPCR)

The expression of epithelial and fibroblast markers was determined in established cell lines using qPCR. Briefly, RNA was extracted from cell lines using the Trizol reagent (Invitrogen, USA) according to the manufacturer’s instructions. The extracted RNA was evaluated for RNA concentration, A260/A280 and A260/230 values using Nanophotometer (Implen, USA). Acceptable A260/280 was 1.8–2.2 and A260/230 was 2.0-2.2. If not immediately used, the RNA was stored at −80°C until analysis. cDNA conversion was achieved using the RevertAid First Strand cDNA synthesis kit (Thermo Fisher, USA) according to the manufacturer’s instructions.

Quantitative PCR reaction was performed with the EvaGreen qPCR Mix Plus (Solis Biodyne, Estonia). The run method used was as follows :

Holding Stage – 95°C, 15 minutes

Cycling – Denaturation 95°C, 15 seconds

Annealing 60°C, 1 minute

Elongation 72°C, 20 seconds

Gene expression level was quantified by the comparative C_T_ method. The difference between the C_T_ values (Δ C_T_) of the gene of interest and the housekeeping gene was calculated for each experimental sample. The housekeeping gene used was 18s rRNA. The expressions of epithelial markers cytokeratin 8 (CK8) and E-cadherin, as well as fibroblast markers alpha smooth muscle actin (α-SMA), fibroblast activation protein (FAP) and vimentin were evaluated. Vimentin was expected to be expressed in ccRCC epithelial cells as well. Primer sequences are shown in Table 2. Primer sequences were obtained from PrimerBank, a validated public online resource for PCR primers provided by collaboration between Massachusetts General Hospital and Harvard Medical School. Gene expression of established cell lines were compared to the equivalent mRNA levels found in a representative ATCC RCC epithelial cell line (ACHN). All experiments were carried out in triplicates.

**Table 2.**
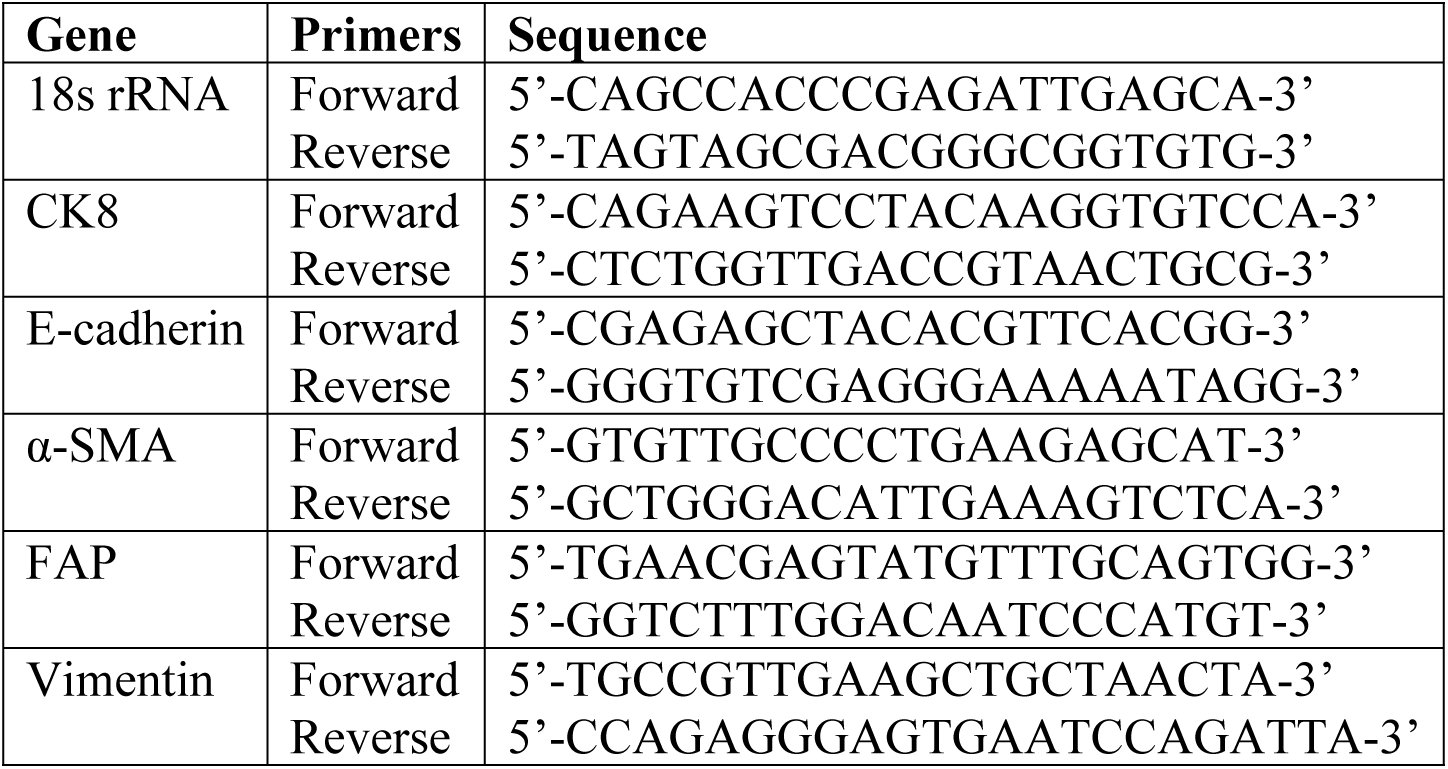
Primer sequences for qPCR of epithelial and fibroblast marker expressions in cell lines.

## Results

### Establishment of renal tumour and normal kidney cell lines

After optimizing the establishment method, 11 cell lines spontaneously immortalised out of 51 trials (from different patients). Therefore, the spontaneous immortalisation rate of tumour epithelial cells was 21.6% with the optimized protocol. Cells were considered immortalised if they could be passaged beyond the 10^th^ passage. Most cells senesced and stopped proliferating after 3-5 passages. Out of the spontaneously immortalised cell lines, 7 were clear cell RCC (ccRCC) variants, 2 were ccRCC with sarcomatoid transformation, one was mostly undifferentiated RCC with some papillary features and one was non-RCC Ewing’s sarcoma. Each cell line had distinctive morphology ranging from polygonal epithelial to spindle shaped and elongated cells (Fig 2). Nine immortalised cell lines were from patients with stage T2 tumours and above, whereas two were stage T1b with metastasis. In total, 6/11 (54.5%) immortalised cell lines were from patients who had metastasis at presentation. The cell lines were from tumours with grade 2 and above except for two with grade 1 and metastasis (both T1b tumours). Fig 3 displays examples of cell lines of various subtypes which grew successfully and their different cell morphologies under the inverted microscope.

**Fig 2.**
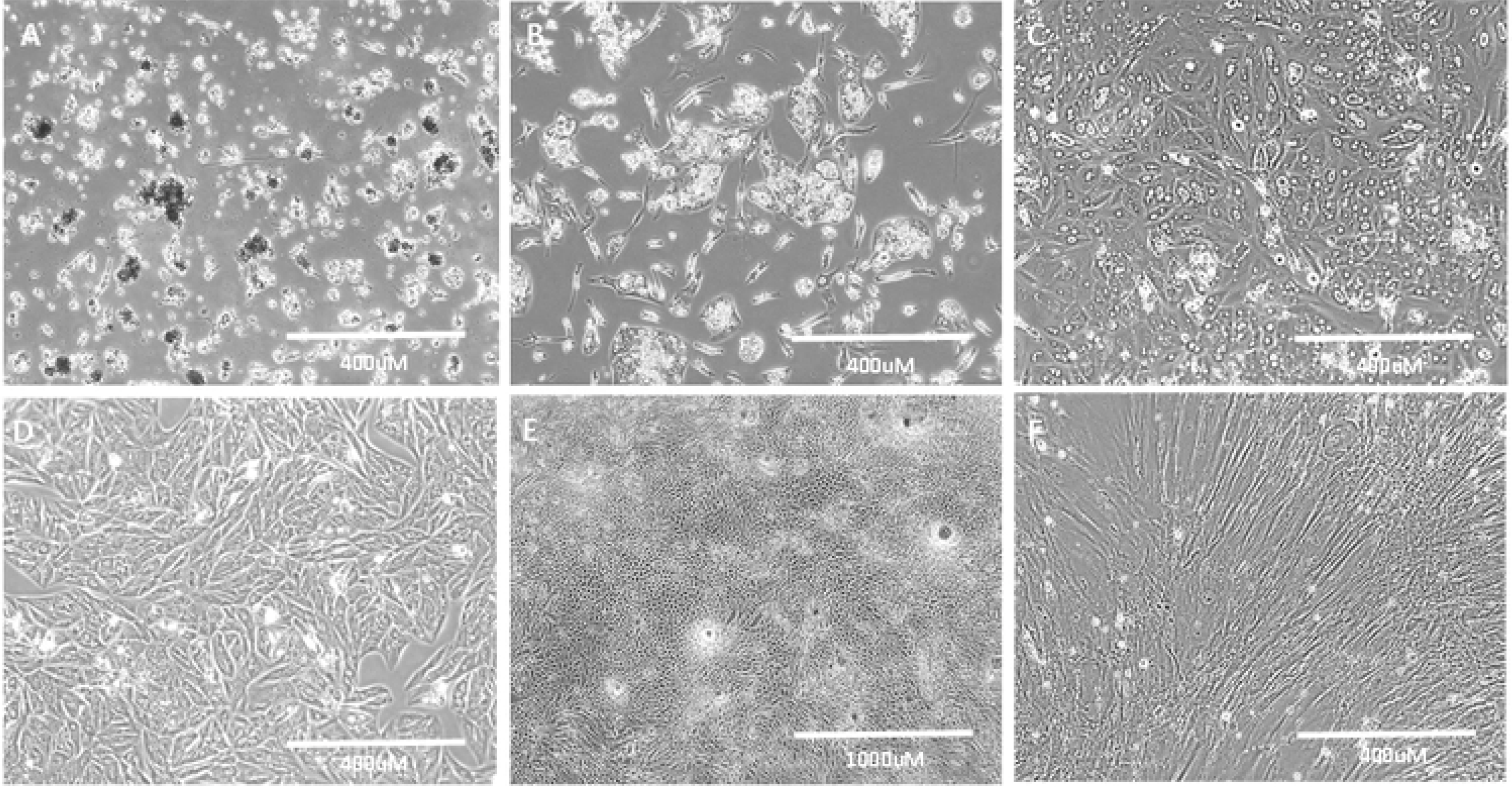
Growth characteristics of culture cells. (A) Day 1 : Attachment of clumps of cells onto tissue culture flask. (B) Day 5 : Cells flatten out and start to proliferate. (C) Day 14 : Cells achieving confluency with lipid/glycogen granules visible in cells. This could be seen in some clear cell RCC primary cells because this subtype is rich in lipid/glycogen. (D) Another ccRCC cell line at confluency which exhibits different morphology from (C) (E) Dome formation is visible in confluent normal kidney culture, which is characteristic of proximal tubule culture (F) Long spindly elongated cells of ccRCC fibroblast All taken at 100x magnification except (E) which was taken at 40x magnification

**Fig 3.**
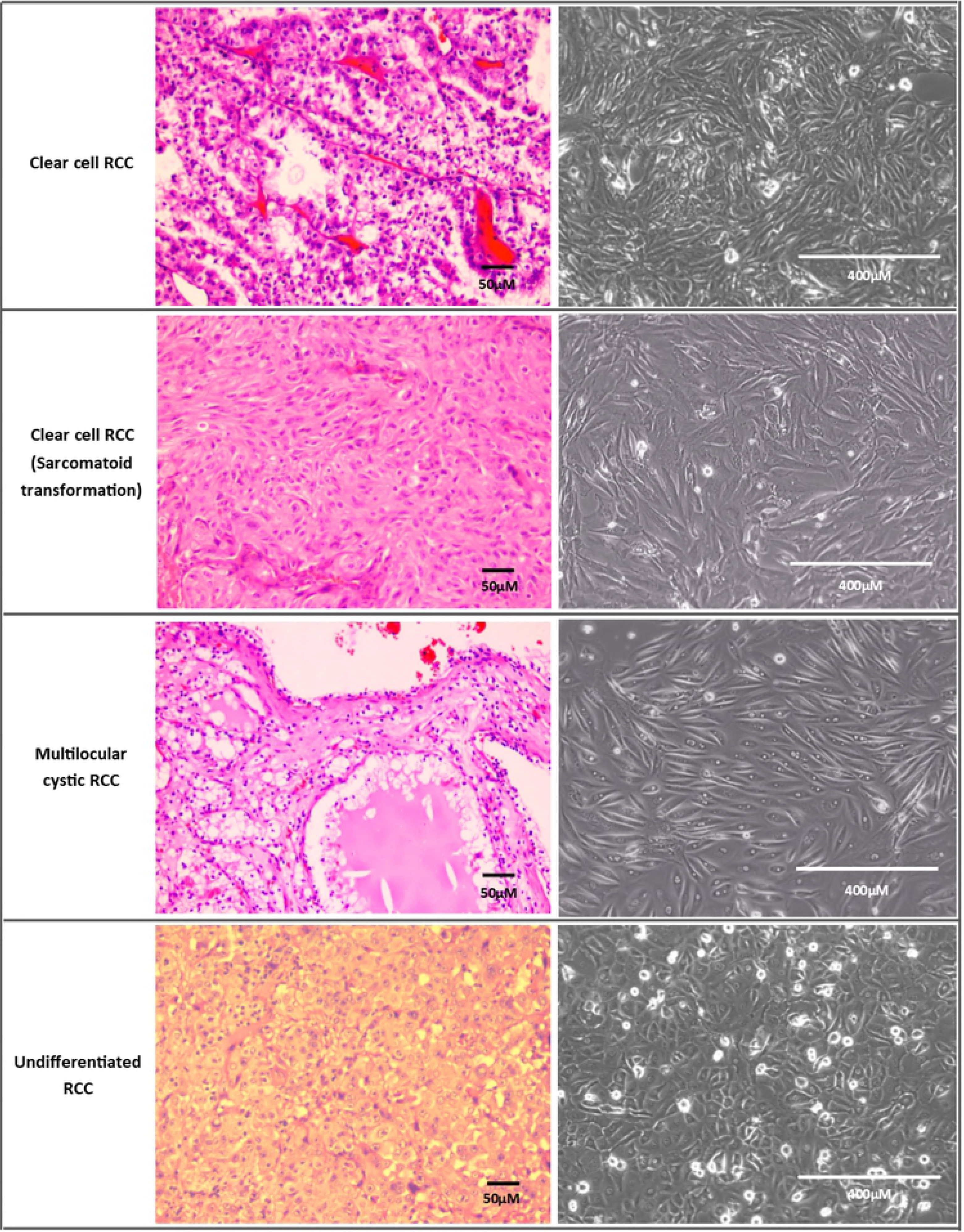
The morphology of cell lines at confluency (<passage 10) and their tumour counterparts (H&E staining) of various subtypes.

Normal epithelial kidney cortex cells (proximal tubule cells) were relatively easy to grow and proliferated at a faster rate than the tumour epithelial cells. After the first passage, tumour epithelial cells usually reached confluency in 3-15 days while normal epithelial kidney cortex cells reached confluency in 2-5 days with a 1:2 split ratio for passaging. However, none of the normal kidney cortex cells achieved spontaneous immortalisation.

Out of 18 trials of growing RCC cancer associated fibroblast cells, none has immortalised at the moment. Eight fibroblast cell cultures were successfully passaged once, with one cell culture proliferating up to nine passages before senescence. The rest either grew too slowly and could not reach confluency to be passaged, were overtaken by epithelial cells or the explant did not attach well. Compared to RCC epithelial cells, RCC fibroblasts were harder to grow and contamination of fibroblast cells in RCC epithelial lines was rarely an issue. Morphologically, RCC epithelial and fibroblast cells were distinguishable (Fig 2).

### Immunofluorescence characterisation of cell lines

RCC epithelial cells stained strongly positive for the epithelial marker, pan-cytokeratin and negative for the fibroblast marker, α-SMA (Fig 4). Fibroblast cells stained strongly for α-SMA and negative for pan-cytokeratin. Normal kidney cortex cells (proximal tubule cells) stained positive for pan-cytokeratin and AQP-1 (proximal tubular epithelial cell marker) (Fig 4) while staining negative or weakly for α-SMA and the distal tubule marker THP (images not shown).

**Fig 4.**
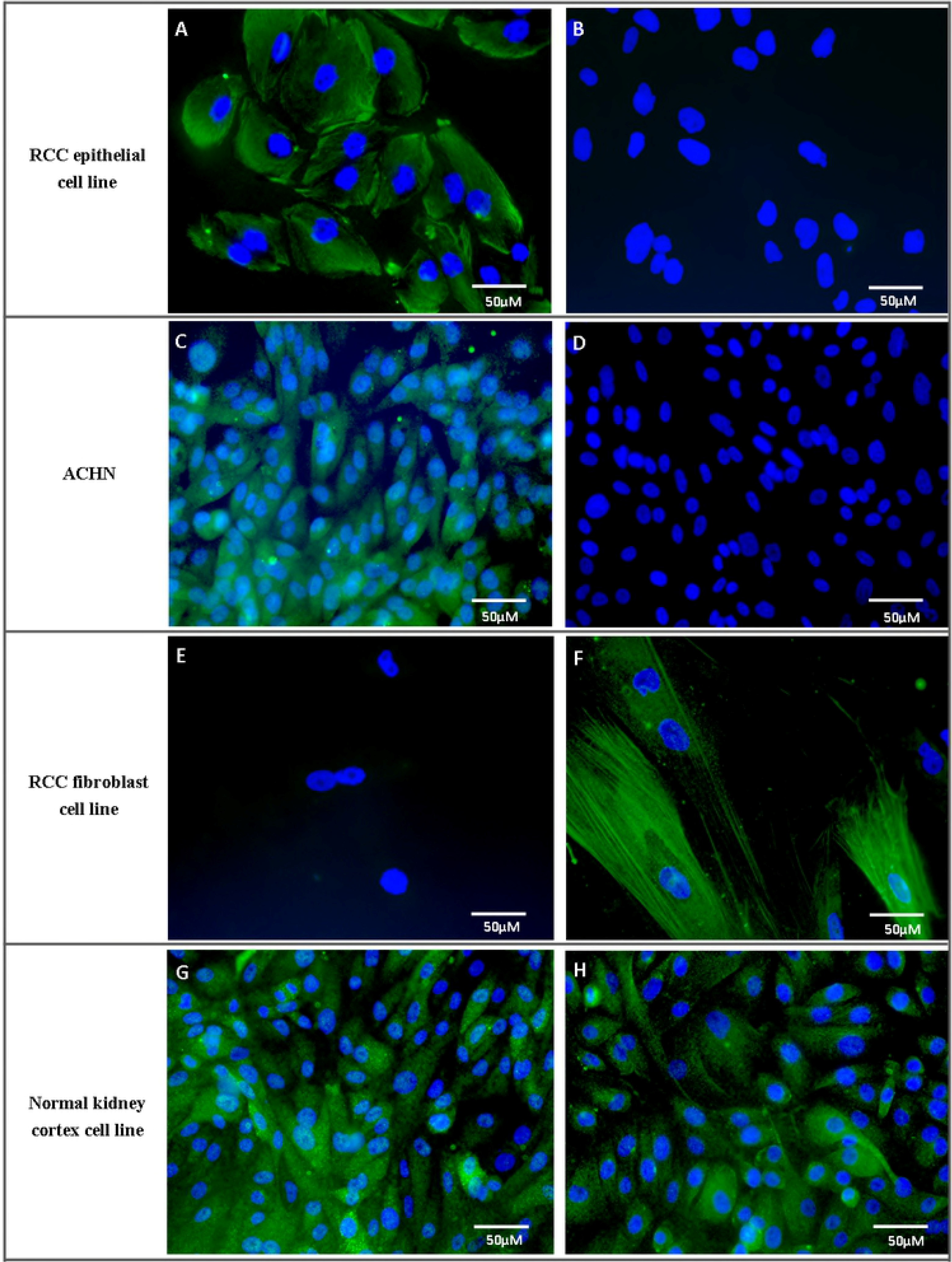
Examples of DAPI and Alexa fluor 488 staining of established RCC epithelial, RCC fibroblast and normal kidney cortex cell lines. Commercial ACHN cell line was used as a comparison. A, C, E and G depict staining for pan-cytokeratin while B, D and F are staining for α-SMA. H shows staining for AQP-1 in normal kidney cortex cells.

### Quantitative PCR characterisation of cell lines

Quantitative PCR analysis revealed higher epithelial marker expressions in these selected examples of RCC epithelial cell lines compared to fibroblast cells (Fig 5). These epithelial cell lines were designated UMRCC1 onwards. UMRCC1, UMRCC2, UMRCC6 and UMRCC10 showed higher epithelial marker (CK8 and E-cadherin) and lower fibroblast marker (α-SMA) expressions compared to Fibroblast 1 and 2. The RCC epithelial cell lines had lower FAP expression compared to the fibroblast cells except UMRCC6 which was established from a ccRCC tumour with sarcomatoid transformation. All cell lines have mixed vimentin (fibroblast marker) expression as ccRCC epithelial cells are known to express this protein.

**Fig 5.**
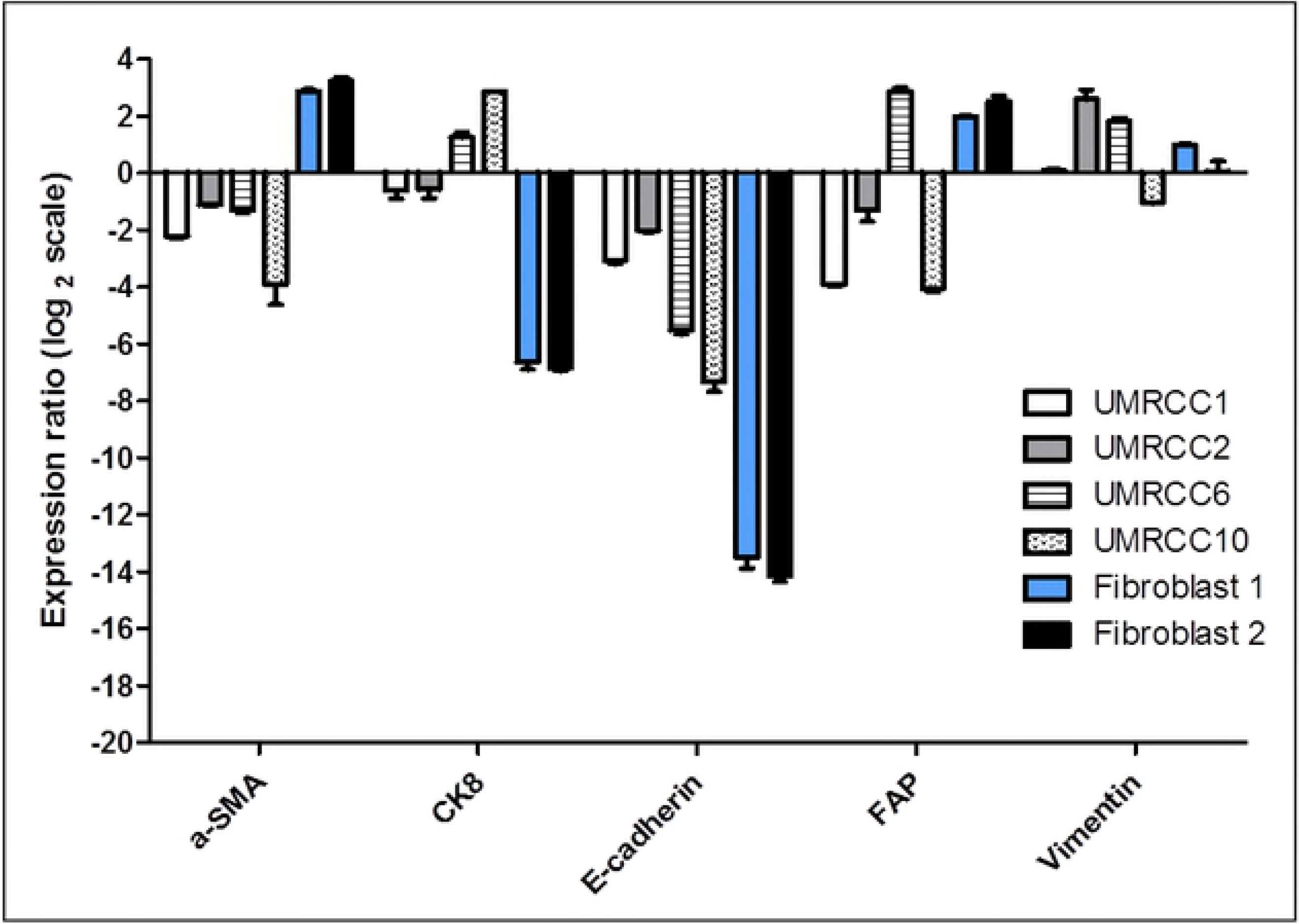
Epithelial and fibroblast marker expressions of selected RCC epithelial cell lines (UMRCC1, UMRCC2, UMRCC6 and UMRCC10) and RCC fibroblast cells (Fibroblast 1 and 2). Data shown are representative of triplicate experiments. The expression ratio of the genes in the newly established cell lines were compared to the equivalent mRNA levels found in a representative ATCC RCC epithelial cell line (ACHN).

## Discussion

Epithelial RCC cell lines were established from the primary tumour tissue of RCC patients with a spontaneous immortalisation rate of 21.6% using this protocol. This was slightly higher than the 12.7% rate obtained by Ebert et al. (1990), which reported the spontaneous immortalisation rate of RCC cells [9]. In this study, immortalised cell lines were from tumours with more clinically aggressive characteristics such as larger tumours (stage T2 and above), higher grade (grade 2 and above) or has metastasized. The established cell lines were confirmed to be epithelial cells with higher expressions of epithelial markers such as pan-cytokeratin, CK8 and E-cadherin [7,14]. These cells also exhibit lower expression of fibroblast marker α-SMA [15]. Interestingly, UMRCC6, a RCC cell line with sarcomatoid differentiation, had higher FAP expression compared to the other epithelial cell lines evaluated. FAP is marker of activated fibroblast and is also expressed by cancer associated fibroblasts. In primary RCC tumours, FAP is expressed on stromal fibroblasts and is shown to be associated with tumour aggressiveness, including sarcomatoid transformation [16]. Morphologically, UMRCC6 cells were slightly spindle shaped and expressed high epithelial markers.

Using this protocol, kidney tumour cell lines of various subtypes were successfully established, including uncommon types like ccRCC with sarcomatoid transformation, largely undifferentiated RCC and Ewing’s sarcoma, which is a non-RCC kidney tumour. A multilocular cystic RCC cell line was successfully grown for three passages, but did not achieve spontaneous immortalization. Most commercially available cell lines are of the common subtype, ccRCC, such as Caki-1, Caki-2, 769-P and 786-0, while the subtypes of ACHN and A-498 cell lines were unclear [17]. Hence, these newly established cell lines could provide a valuable resource as in-vitro models for rare kidney tumour subtypes. To the authors’ knowledge, there are no commercial RCC with sarcomatoid features or Ewing’s sarcoma of the kidney cell lines which are easily available.

Normal kidney cell lines established from the same patients can be used as controls for invitro experiments, as they represent the non-malignant counterpart of the tumour cell line. In addition to established cell lines (≥ passage 1), cryo-preserved primary culture (passage 0) is available for studies with the likelihood that they retain the molecular profile of the corresponding tissue [11]. The above reasons emphasize the advantage of establishing cell lines at research centres if patient samples are accessible.

Several protocols for RCC and kidney cell line establishment have been reported, utilizing various techniques such as the gradient centrifugation technique, enzymatic digestion, or explant method [7-12]. Based on the trials during this protocol optimization, the growth of the epithelial cells before the first passage for the explant method was slower compared to using enzymatic tissue digestion. Therefore, the epithelial protocol described here entails enzymatic digestion followed by cell sieving. Contamination with fibroblast cells was seldom an issue in this study as RCC tumour associated fibroblast cells were more difficult and slower to grow than the epithelial cells. Fibroblast cells were more successfully grown using the explant method, similarly described by previous groups [13,18]. Due to the limited publications on cancer associated fibroblasts in RCC, the optimization of the fibroblast establishment protocol or technique could be pursued further. This is because cancer associated fibroblasts are known to interact with cancer cells to promote tumour growth and progression [19].

Compared to previously reported protocols, additional antibiotics and washing steps were added as a precaution to prevent bacterial and mycoplasma growth as primary cultures from human tissues can suffer from contamination issues [7-12,20,21]. Using this protocol, coating of culture flasks is not required and washed tumour/normal kidney tissue can be left overnight in culture medium at 4°C before further tissue processing with good success rate. This allows for more convenient tissue processing of specimens collected from operation cases which are carried out in the evenings or at night.

## Conclusions

In summary, the protocol described in this paper allows for simultaneous establishment of RCC, normal kidney and RCC associated fibroblast cell lines or cultures from nephrectomy specimens with a good success rate of spontaneous immortalization for RCC epithelial cell lines. Tissue location selection from the surgical specimen is important and morphological, immunofluorescence and qPCR characterization can be carried out to determine the epithelial or fibroblastic origin of the cells. Further characterization via mutation analysis can be performed next to determine the genetic mutational or molecular features of the cell lines.

## Acknowledgements

This study was funded by Ministry of Education MOSTI grant 01-02-03-SF0919 and University of Malaya PPP Grant PG140-2015B

